# Contributions of action potentials to scalp EEG: theory and biophysical simulations

**DOI:** 10.1101/2024.05.28.596262

**Authors:** Niklas Brake, Anmar Khadra

## Abstract

Differences in the apparent 1/f component of neural power spectra require correction depending on the underlying neural mechanisms, which remain incompletely understood. Past studies suggest that neuronal spiking produces broadband signals and shapes the spectral trend of invasive macroscopic recordings, but it is unclear to what extent action potentials (APs) influence scalp EEG. Here, we combined biophysical simulations with statistical modelling to examine the amplitude and spectral content of scalp potentials generated by the electric fields from spiking activity. We found that under physiological conditions, synchronized aperiodic spiking can account for at most 1% of the spectral density observed in EEG recordings, suggesting that the EEG spectral trend reflects only external noise at high frequencies. Indeed, by analyzing previously published data from pharmacologically paralyzed subjects, we confirmed that the EEG spectral trend is entirely explained by synaptic timescales and electromyogram contamination. We also investigated rhythmic EEG generation, finding that APs can generate narrowband power between approximately 60 and 600 Hz, thus reaching frequencies much faster than the timescales of excitatory synaptic currents. Our results imply that different spectral detrending strategies are required for high frequency oscillations compared to slower synaptically generated EEG rhythms.

## Introduction

Understanding the neural mechanisms underlying EEG generation is important for inferring changes in brain state, as well as developing methods to filter out irrelevant signals. Towards this latter aim, recent work has focused on characterizing the neural basis of broadband EEG signals and defining when and how EEG spectra need to be detrended^1–3^. Studies into the neural basis of broadband EEG have primarily focused on synaptic filtering^3–6^ and low frequency, aperiodic network fluctuations^3, 7, 8^. However, in addition to synaptic contributions, the spectral trend observed in invasive, large-scale neural recordings, such as the local field potential (LFP)^9–11^ and intracranial EEG (iEEG)^12, 13^ including electrocorticography (ECoG)^14–17^, is believed to reflect broadband contributions from spiking activity^9, 10, 18^, especially at frequencies above *∼*60 Hz, the so-called high gamma range. Such high frequency broadband contributions are thought to be important for determining the slope of the 1/*f* spectral trend^19^.

In comparison to invasive recording techniques, the majority of the unprocessed EEG signal above 30 Hz reflects muscle activity^20–23^. Moreover, EEG is thought to be incapable of measuring APs because they are believed to be too brief and unsynchronized^24, 25^. Nonetheless, when muscle artifacts are corrected for, EEG recordings have displayed transients in the high gamma range^26–28^, similar to those observed in LFP and iEEG recordings. If such high frequency transients are indeed generated by synchronized APs, it would hold significant implication for interpreting spectral peaks and correcting for the EEG spectral trend. Interestingly, a recent biophysical modelling study showed that APs account for almost 20% of the amplitude of single-neuron dipoles, and concluded that APs can contribute significantly to EEG rhythms^29^. However, a systemic investigation into the ability of APs to produce detectable scalp potentials has not been undertaken. Additionally, the potential contribution of APs to aperiodic EEG signals and the overall spectral trend has not been explored.

In this study, we aim to address this gap by employing a quantitative approach that explores AP-generated EEG signals, a type of signal that we refer to hereafter as apEEG for brevity. To begin, we employ a combination of biophysical simulations and statistical modelling to examine the impact of single neuron properties and spike synchrony on the amplitude and spectral features of apEEG signals. Using these results, we evaluate whether apEEG can exhibit experimentally-measurable narrowband and broadband high gamma power. Our results have implications for interpreting high frequency EEG rhythms and for designing practical methods for spectral detrending.

## Results

### Unitary AP response of single-neuron dipoles is approximately linear

The contributions of an individual neuron to the electric potential measured by a distant electrode can be modelled by a single dipole vector that varies with time^30, 31^. This case applies well to EEG signals due to the distance between the brain and scalp electrodes. To understand how APs contribute to EEG, we therefore first sought to characterize the contributions of APs to their respective neurons’ dipoles. We simulated neuron models with detailed morphologies and distributed passive and voltage-gated ion channels on the soma, axon initial segment, and dendrites (**Fig. 1A**). To induce spiking, we bombarded the dendrites of this active model with background synaptic input, whereas to block spiking in the presence of such synaptic inputs, we set the conductance of voltage-gated sodium channels in the soma and axon initial segment to zero, obtaining a passive model that allowed us to characterize the dipole generated in the absence of firing (**Fig. 1B**). By subtracting the active and passive simulation results and thereafter taking the spike-triggered average of the single-neuron dipole, we estimated the unitary AP response of the electric field (**Fig. 1C**).

**Fig 1.**
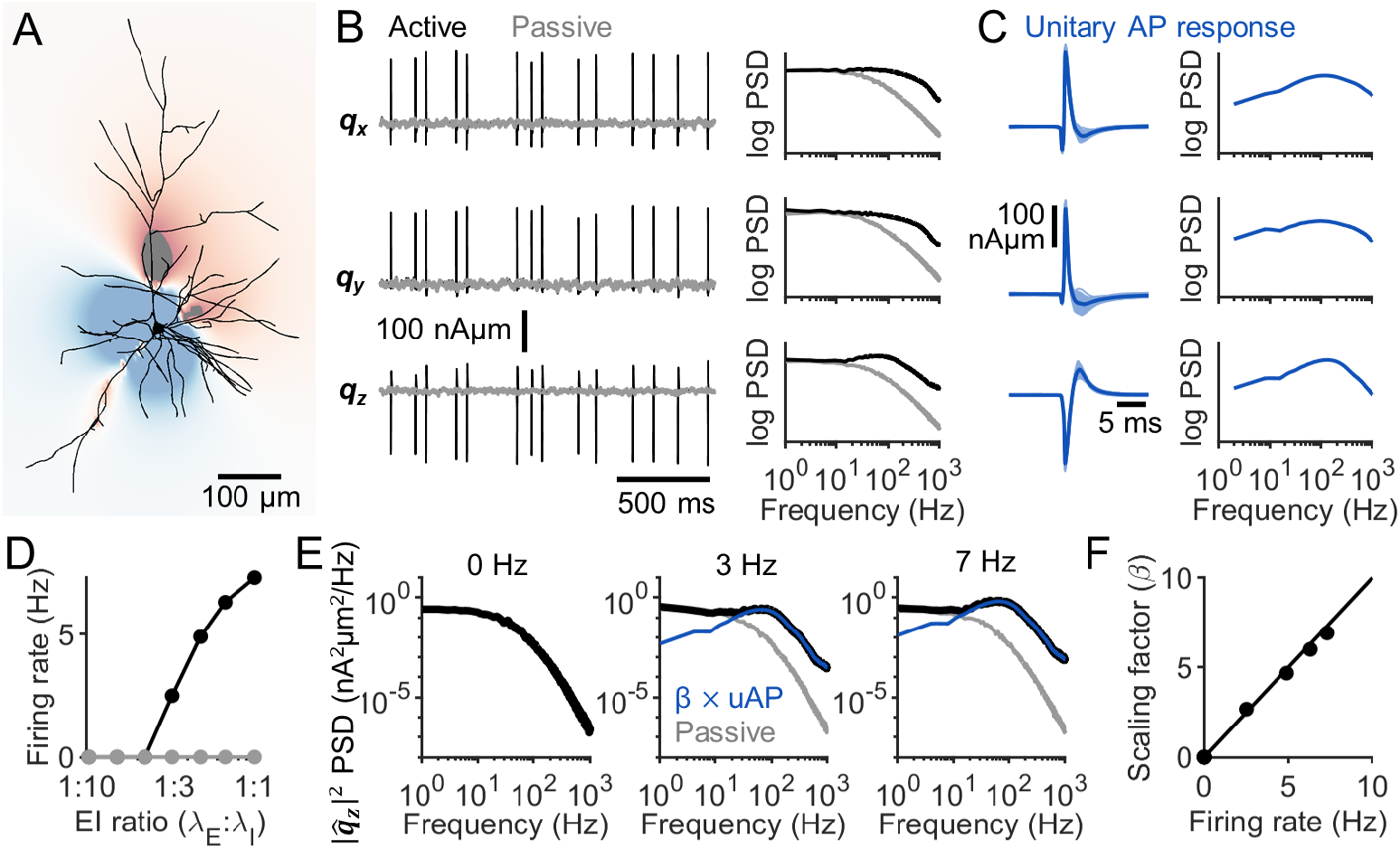
Calculating the unitary AP response. (**A**) The extracellular electric field generated by a neuron is shown at the peak of an AP. (**B**) Left: The single-neuron dipole, ***q***, associated with the neuron in panel A for the active model (black) and passive model with sodium channels removed (grey). The x, y, and z components of the vector are plotted from top to bottom. Notice the correspondence in the subthreshold fluctuations between the two sets of simulations. Right: The power spectrum of each dipole component trace on the left for the active (black) and passive (gray) model. (**C**) Left: The difference between the single-neuron dipoles calculated with the active and passive model aligned to each AP (light blue), along with the spike-triggered average (dark blue) which defines the unitary AP response. x, y, and z components are shown from top to bottom. Right: the power spectrum of the unitary AP response. (**D**) The firing rate of the active (black) and passive (gray) model as a function of E:I ratio, defined as the ratio between the rate of excitatory synapse activation, *λ*_*E*_, to that of inhibitory synapse activation, *λ*_*I*_. (**E**) The power spectrum of the z component of the single-neuron dipole (black) at three different firing frequencies: 0 Hz (left), 3 Hz (middle) and 7 Hz(right), along with the spectra of the passive models (gray) and the spectra of the unitary AP response (blue) shown previously in panel C. Notice how the unitary AP spectrum matches, up to a scaling factor (*β*), the single-neuron dipole spectrum at high frequency. (**F**) The scaling factor for the unitary AP spectrum that fits the single-neuron dipole spectrum, plotted as a function of the firing rate (black dots). These data points almost align perfectly with the unity line (black line). The x and y components of the dipole vector show the same behaviour (**Fig. S1**).

The ensemble electric field is equal to the linear summation of those generated by each individual neuron in the brain^24, 32^. However, the electric fields generated by individual neurons are not in general linear; sublinear and supralinear interactions among synaptic currents prevent this^33^. Nonetheless, one might hypothesize the contributions of APs to be linear. In this case, the spectrum of the single-neuron dipole, *S*(*f*) would be proportional to the energy spectrum of the unitary AP response, *S*_*ap*_(*f*), satisfying the equation

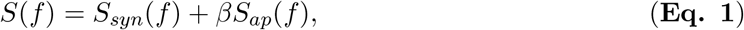

where *S*_*syn*_(*f*) is the power spectrum of the synaptic contributions, and *β* is a scaling factor that should be equal to the cell’s firing rate, as we demonstrate below. To test the accuracy of this simplified model, we calculated single-neuron dipoles while varying the firing rate of the neuron by altering the ratio of excitatory to inhibitory input (**Fig. 1D**). We estimated *S*_*syn*_ for each EI ratio by considering the passive model in which the sodium channel conductance was set to zero. Meanwhile, *S*_*ap*_ was defined as the energy spectrum of the unitary AP response calculated at low firing rates (see Methods). By fitting the power spectrum of the single-neuron dipole at each EI ratio with **Eq. 1**, we estimated the scaling factor *β* and showed that it closely matches the firing rate (**Fig. 1E, F**).

We performed this analysis on biophysical models of 68 representative neuron classes^34^ (**Table S1**). Across all models, the unitary AP scaling factor *β* closely followed the firing rate (**Fig. 2A**). However, as the firing rate increased, the accuracy of the linear approximation decreased (**Fig. 2B**). This was because the unitary AP responses were less representative of the AP responses occurring at high frequencies. Nonetheless, we found that the spectral profile of the AP-generated signal was nearly identical to that predicted by the linear model up to firing rates of approximately 80 Hz (**Fig. 2C**). We concluded that the amplitude of AP responses are captured well by a linear model, but that the precise spectral properties of these responses may be slightly different than those predicted by a fully linear model during sustained high frequency firing above 80 Hz. However, considering that the average firing rates of active neurons typically fall below 60 Hz^35–38^, we deemed this simplification of AP signals to be acceptable.

**Fig 2.**
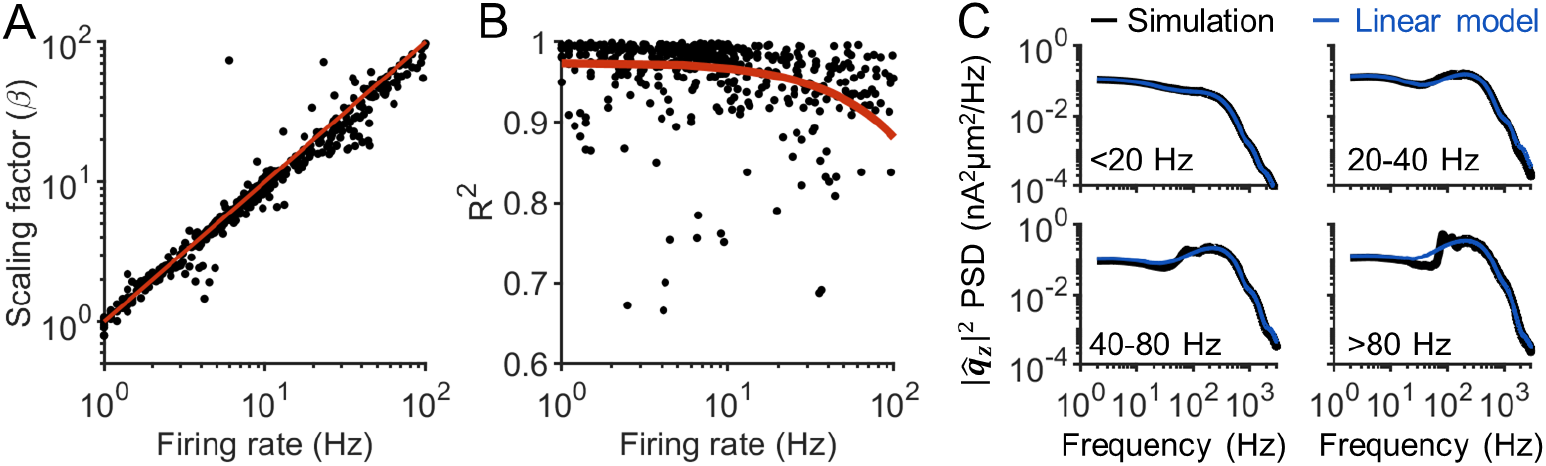
AP contributions to single-neuron dipoles are linear with firing rate. (**A**) Fitted unitary AP scaling factor (*β*; see **Eq. 1**) plotted against firing rate for 68 neuron models covering the 55 neuron classes identified by Markram et al.^34^ (**Table S1**). These data points align almost perfectly with the unity line (red line). (**B**) The *R*^2^ value obtained from fitting a linear model (**Eq. 1**) to the spectra of the AP dipole response in each simulation. Notice how the line of best fit (red line) shows that **Eq. 1** gets less accurate at high firing rate. (**C**) The spectra of the z component of the single-neuron dipoles (black), averaged across all simulations with firing rates in the specified ranges, compared to the spectra predicted from the linear model (blue). For firing rates less than 80 Hz, the linear model produces spectra nearly identical to the simulations. At firing rates above 80 Hz, there is a slight departure is spectral density around 100 Hz. The same results were obtained with the x and y components of the dipole (**Fig. S1**).

### A linear model for the spectrum of AP electric fields

Based on the foregoing linearity assumption outlined in **Eq. 1**, we derived a general equation for the ensemble electric fields generated by APs. In general, the potential between two electrodes can be calculated from their lead field^32^, ***ν***(***x***), which describes the sensitivity of the measured potential with respect to a unit dipole vector at the spatial point ***x***. Using this formalism, we can write the potential generated by *N* neurons as

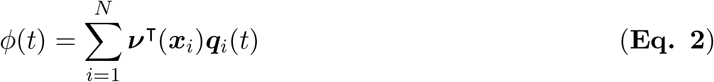

where ***q***_*i*_ is the single-neuron dipole of neuron *i*, located at coordinate ***x***_*i*_ in the brain. This equation leads to the following power spectrum for the ensemble signal

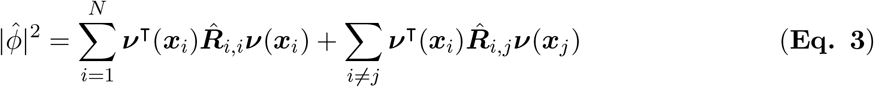

where ***R***_*i,i*_(*τ*) is the auto-correlation matrix of the single-neuron dipole for neuron *i*, ***R***_*i,j*_(*τ*) is the cross-correlation matrix for two neurons *i* and *j*, and 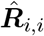 and 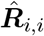 denote their Fourier transforms, respectively.

We now make use of the linearity result from the previous section by describing ***q***_*i*_ as a linear filter, 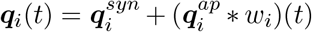, where the vector 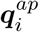 is the unitary AP response of neuron *i, w*_*i*_ is a point process describing the spike times of the neuron, and 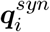 is the synaptic component of the single-neuron dipole, assumed to be statistically independent of 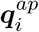 (**Eq. 1**; see also Discussion). The AP component of the ensemble potential is therefore

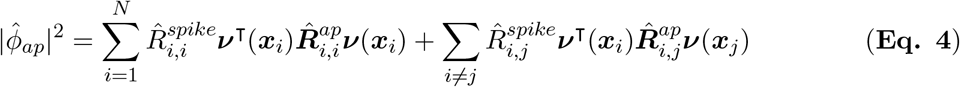

where 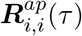 is the auto-correlation matrix of the unitary AP response for neuron *i*, 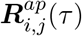 is the cross-correlation matrix for two neurons *i* and *j*, 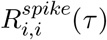 is the spike train auto-correlation of neuron *i*, 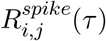 is the spike train cross-correlation of neurons *i* and *j*, and 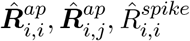 and 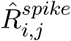 denote their Fourier transforms, respectively.

We estimated the average auto- and cross-correlations between unitary AP responses based on biophysical simulations of all 1035 neuron models generated by the Blue Brain project^34^. As expected, when the neurons fired APs with zero lag, their dipoles exhibited strong cross-correlations along the apical-basal axes of their respective neurons (**Fig. S2**). Interestingly, these calculations also revealed significant cross-correlations between the dipoles’ apical-basal component and their azimuthal components (**Fig. S2**). This observation suggests that even neurons that are not aligned in parallel may still generate coherent electric fields during synchronous firing, thus further boosting the signals generated by populations of AP responses.

### Magnitude of apEEG depends on dendrite asymmetry

Using the above linear model, we sought to calculate the potential measured at an EEG electrode generated by APs under various conditions. To begin, we investigated the simple case where neurons fire according to uncorrelated Poisson spike trains. In this case, 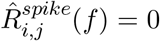 and 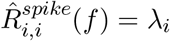, although for simplicity we assumed a single average *λ* for all neurons. To estimate the solution to this equation, we divided our neuron models into the 55 different morphology classes defined by Markram et al.^34^ (**Table S2**). Under this scenario, the apEEG spectrum was calculated as

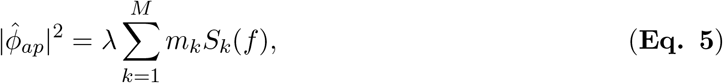

where *S*_*k*_ is equal to the expected EEG spectrum generated by a neuron of class *k* firing a single AP, and *m*_*k*_ is the number of neurons that fall into each class. *S*_*k*_ was calculated by averaging the EEG spectra generated by simulating many neurons of class *k* and placing them at each of the 75,000 cortical locations available in the New York Head model^39^ (**Fig. 3A**). This is identical to the procedure used previously to calculate the unitary spectrum for the synaptic component of the EEG^3^. To then calculate the ensemble apEEG spectrum, the number of neurons in each class, *m*_*k*_, was calculated by multiplying the estimated abundance of each cell type^34^ (**Fig. 3B**) by the total number of neurons in the cortex, which we took to be 16 billion^40^.

**Fig 3.**
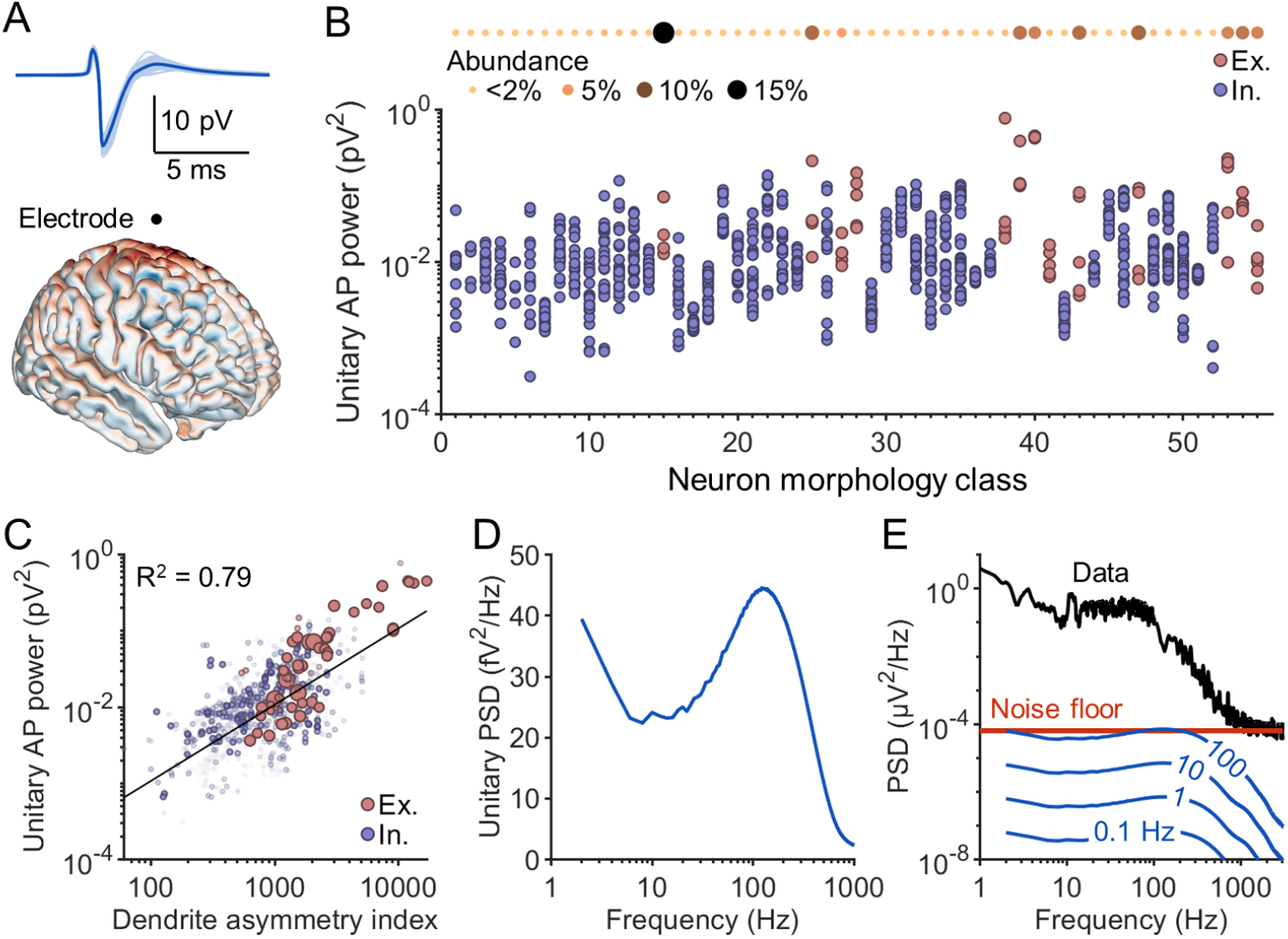
Diverse cell types’ unitary AP contributions to scalp EEG. (**A**) An example unitary apEEG response (top) computed by placing a simulated single-neuron dipole at a random cortical location in the New York Head model (bottom) and calculating the signal at the Cz electrode site. (**B**) The unitary apEEG power, averaged across simulations of all possible neuron locations in the cortex, for each of the 1035 neuron models, split into various morphology classes; a description of each morphology class is provided in **Table S2**. Excitatory neurons are shown in red and inhibitory neurons in blue. The relative abundance of each morphology type is shown at the top of the panel. (**C**) The location-averaged unitary apEEG power of each neuron model plotted against the neuron’s dendrite asymmetry index (**Eq. 8**, see Methods). The size and opacity of each point is directly proportional to the neuron’s relative abundance in the brain. Black line: line of best fit. (**D**) The expected unitary apEEG spectrum, averaged over all neuron models and weighted by the relative abundance of each neuron type. (**E**) The power spectrum of EEG collected by Scheer et al.^41^ (black) and the associated noise floor (red). Blue lines: the simulated apEEG spectrum generated by the entire brain firing asynchronously at various frequencies.

Among neuron classes, the average power of the unitary apEEG response varied by almost two orders of magnitude (**Fig. 3B**). Excitatory pyramidal cells tended to generate larger amplitude apEEG signals than inhibitory neurons, as expected^29^. However, certain inhibitory neurons also generated surprisingly large amplitude signals (**Fig. 3B**). Whereas the average excitatory neuron generated a unitary apEEG response with an energy of *∼*0.09 pV^2^, the average inhibitory neuron generated signals of *∼*0.02 pV^2^. Because pyramidal neurons are thought to dominate EEG signals due to their polarized dendrite morphology, we hypothesized that many interneurons have significant asymmetries in their dendritic arbours. To test this, we defined a dendrite asymmetry index (**Eq. 8**; see Methods) and evaluated the predictive power of this measure on apEEG signal strength. Consistent with our hypothesis, the unitary apEEG power for each neuron was strongly correlated with its dendrite asymmetry index (**Fig. 3C**). While in general excitatory neurons exhibited more dendrite asymmetry, many interneuron dendrites displayed equal or greater asymmetry (**Fig. 3C**). This result demonstrates that interneuron spikes can generate large electric fields, commensurate with those of many excitatory neurons.

Interestingly, the expected unitary apEEG spectrum revealed both low pass and bandpass properties (**Fig. 3D**). The bandpass property, which is reflected in the peak in the power spectrum around 100 Hz, arises from the fast temporal dynamics of the up and downstroke of the AP waveform. The low-pass filtering properties are evident in the low frequency power below 10 Hz. This power was disproportionately contributed by certain neuron classes which exhibited significant, slow after-hyperpolarizations that often took tens to hundreds of milliseconds to return to baseline (**Fig. S3**).

Finally, we examined the ensemble apEEG spectrum (**Fig. 3D**). Even with an unrealistically high brain-wide firing rate of 100 Hz, the amplitude of the ensemble apEEG signal barely reached the noise floor of high resolution, low noise EEG recordings (**Fig. 3E**). Given the absence of synchrony, these spectra serve as indicators for defining lower bounds on any contributions of APs to scalp EEG. Unsurprisingly, asynchronous firing does not produce detectable apEEG signals.

### Spike synchrony cannot produce high frequency broadband apEEG

We next investigated the effects of spike synchrony on apEEG generation, and turned to the full **Eq. 4**. We used a minimal model for spike synchrony based on two general observations:

1. Spike synchrony is strongest among nearby neurons^42–44^. This was implemented in our model by synchronizing the spike timing of neurons depending on their pairwise distance according to 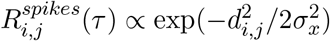, where *d*_*i,j*_ is the Euclidean distance between neurons *i* and *j*, and 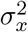 is a parameter that controls the cortical distance over which activity becomes uncorrelated. In accordance with unit recordings in visual cortex^42, 43^, we set 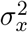 to be 3 mm^2^ (**Fig. 4A**). Although recordings from prefrontal cortex suggest a slightly lower value of around 1 mm^2 45^, differences in the value of *σ*_*x*_ at the scale of millimeters did not have meaningful effects on the results that follow (**Fig. S5**).
2. Even neurons with correlated spiking do not fire at exactly the same time. Therefore, the timescale of correlation was captured by modelling the spike train cross-correlation as a Gaussian function, whose variance, 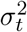, reflects the jitter in spike times^42–44, 46^ (**Fig. 4B**).

**Fig 4.**
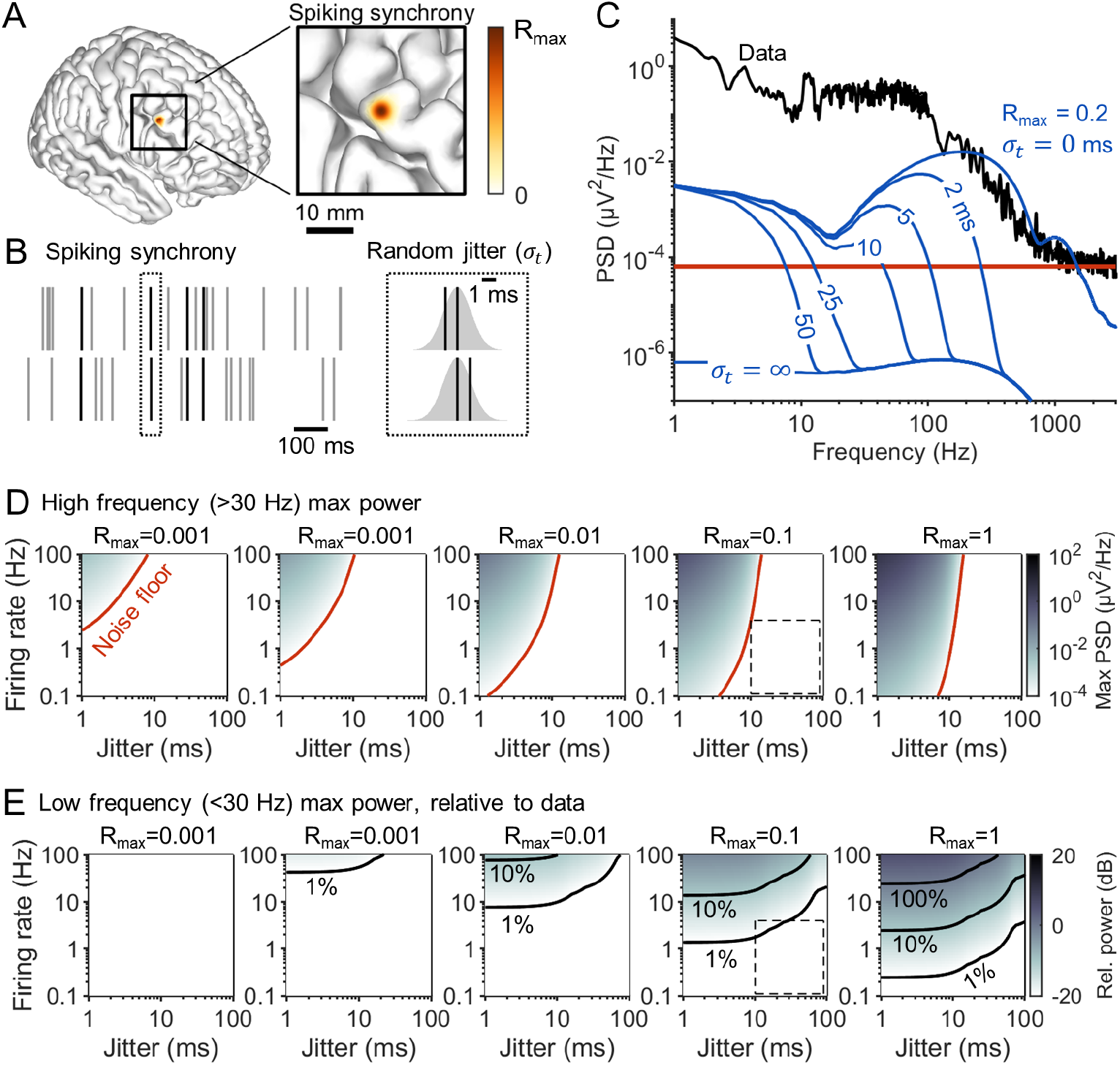
Aperiodic APs cannot generate detectable EEG signals. (**A**) Schematic illustration of the local nature of correlated activity in the model. Neighbouring neurons fire spikes with a correlation of *R*_*max*_, while neuron pairs that are increasingly separated show gradually decreasing correlation. (**B**) Schematic illustrating the timescale of correlation. Correlated neurons have a given fraction of their spikes synchronized (left) with a jitter value drawn from a Gaussian distribution of standard deviation *σ*_*t*_ (right). (**C**) The power spectrum of EEG collected by Scheer et al.^41^ (black) and the example ensemble apEEG spectra of a brain with an average firing rate of 1 Hz and maximal correlation of *R*_*max*_ = 0.2, plotted for various values of *σ*_*t*_ (blue). Red line: The associated noise floor of the experimental EEG spectrum. (**D**) Maximal spectral density above 30 Hz generated by the model for a whole range of firing rates (*λ*) and jitter values (*σ*_*t*_), as well as for various maximal correlation values (*R*_*max*_). Red line: The boundary delineating spectral density above and below the noise floor of the amplifier. Dotted box: the regime of physiologically realistic parameter values (see Methods). (**E**) The maximal power below 30 Hz generated by the model, relative to the experimentally measured spectrum in panel C. Contour lines indicate parameter values where the model generates 1%, 10% and 100% the spectral density of the data. Dotted box: the regime of physiologically realistic parameter values (see Methods).

Together, these two experimental observations give rise to the following equation describing spike synchrony

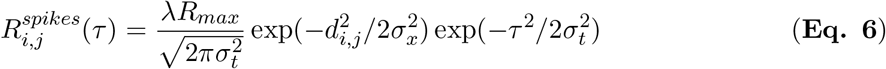

where *R*_*max*_ represents the noise correlation between neurons^46^ and *λ* is the average firing rate of the neurons. According to this model, the dynamics of AP firing and synchrony are both entirely aperiodic, thus allowing us to examine whether apEEG signals can generate aperiodic EEG signals and contribute to the EEG spectral trend.

We estimated the ensemble apEEG spectrum generated by the entire brain using Monte Carlo simulations (see Methods). As a specific example, **Fig. 4C** shows the spectra calculated for *R*_*max*_ = 0.2 and *λ* = 1 Hz. When *σ*_*t*_ = 0, spiking occurs with perfect synchrony, producing an apEEG spectrum that is essentially a scaled version of the average cross-spectrum among unitary AP responses. This spectrum exhibited large amplitude, high frequency broadband EEG signals that would be detectable in EEG recordings. On the other hand, when *σ*_*t*_ = *∞*, the spectrum is identical to the asynchronous case (**Fig. 3**) and would lie far below the noise floor of the experimental EEG spectrum (**Fig. 4C**). For intermediate values, the spectra follow the perfectly synchronous spectrum until a cut-off frequency, determined by *σ*_*t*_, above which the spectrum drops down to the asynchronous spectrum (**Fig. 4C**). This indicates that the timescale of correlation, *σ*_*t*_, is critical in allowing or preventing APs from generating high frequency, broadband EEG signals.

To investigate further, we performed a full sensitivity analysis of model outcomes with respect to the jitter 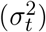, maximal correlation (*R*_*max*_), and firing rate (*λ*). **Figure 4D** illustrates the maximal spectral density produced at frequencies above 30 Hz. The red line indicates where the apEEG crosses the noise floor and the dotted box shows a physiologically reasonable parameter range for *λ, σ*_*t*_, and *R*_*max*_ (see Methods: Determining ranges for parameters). This dotted box is entirely contained below the noise floor of EEG amplifiers, indicating that APs cannot contribute to the high frequency plateau observed in EEG spectra.

At lower frequencies, APs and their after-hyperpolarizations (**Fig. S3**) generated detectable EEG signals even with high jitter values (**Fig. 4C**). To investigate this phenomenon further, we calculated the power generated below 30 Hz as a fraction of that in experimentally recorded EEG, and plotted the maximum obtained power across these frequencies (**Fig. 4E**). For reasonable parameter values, we found that APs could generate maximally *∼*1% of the spectral density seen in recorded EEG signals (**Fig. 4E**). We conclude that while aperiodic APs can generate signals above the detection limit of EEG amplifiers, these signals are dwarfed by the contributions of synaptic currents.

### Excitatory synapses are the only neural sources of spectral trend at high frequency

If APs do not generate detectable aperiodic EEG signals, the EEG spectral trend at higher frequencies should be fully explained by muscle activity^23^ and excitatory synaptic time scales^3, 4^. To test this hypothesis, we analyzed EEG data collected from subjects during peripheral blockade of nicotinic cholinergic receptors^20^, which causes muscle paralysis and removes contamination of EMG signals^20, 22^. In EEG collected from unparalyzed individuals, synaptic timescales were insufficient to explain the high frequency EEG spectral trend especially its plateau (**Fig. 5A**), consistent with previous studies^3^. However, following neuromuscular blockade, this high frequency component of the trend reduced in amplitude in such a way that synaptic timescales could entirely explain the spectral trend (**Fig. 5B**). These results validate our theoretical calculations that APs do not contribute to the EEG spectral trend under baseline conditions, and further validate the role of synaptic timescales in shaping EEG spectra.

**Fig 5.**
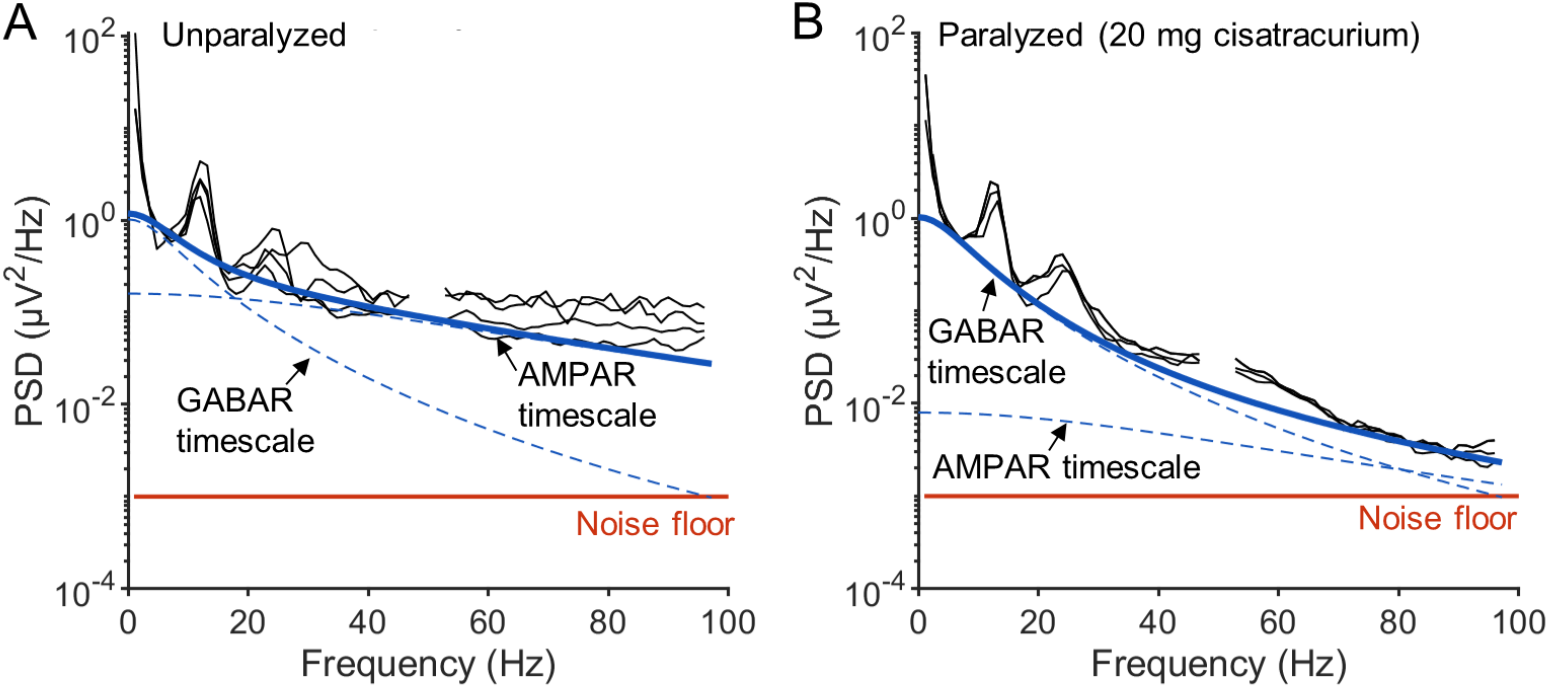
EEG spectral trend is explained entirely by synaptic timescales. (**A**) Spectra of EEG signals collected from unparalyzed subjects by Whitham et al.^20^ (black), fit with **Eq. 9** (solid blue; see Methods). The dashed blue lines indicate the contributions of GABA receptor (GABAR) and AMPA receptor (AMPAR) timescales to the fit. Notice that the fit will never be able capture the high frequency plateau observed in the data. Parameter values: *τ*_*I*1_ = 4 ms, *τ*_*I*2_ = 20 ms, *τ*_*E*1_ = 1 ms, *τ*_*E*2_ = 3 ms, *A*_*I*_ = 3.6, and *A*_*E*_ = 4.6. Red line: Noise floor. (**B**) Same as in A, but with the spectra of EEG signals collected following muscle paralysis and presumed to be free of electromyogram contamination. Parameter values: *τ*_*I*1_ = 4 ms, *τ*_*I*2_ = 20 ms, *τ*_*E*1_ = 1 ms, *τ*_*E*2_ = 3 ms, *A*_*I*_ = 3.6, and *A*_*E*_ = 3.3. Red line: Noise floor.

### Synchronous APs can produce EEG rhythms in gamma range and above

Even if APs do not produce aperiodic EEG signals, it may still be possible that they contribute to EEG rhythms^29^. We therefore used our framework to investigate the consequences of rhythmic synchrony in AP firing activity. We modelled oscillatory synchronization by making the cross-correlation between pairs of neurons a damped sine wave^47, 48^. Mathematically, this means that the spike train cross-correlation is now described by the following equation,

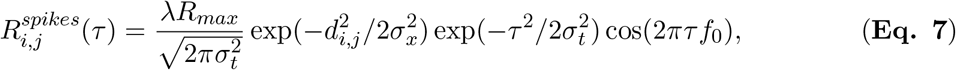

where *f*_0_ is the synchronous rhythm frequency. A representative parameterization of this equation is plotted in **Fig. 6A**. Based on this, we investigated the EEG spectra produced by APs when the frequency of synchronous oscillation, *f*_0_, was systematically varied. The value of *σ*_*t*_ only altered the sharpness of the oscillations spectral peak, but did not affect its amplitude (**Fig. S6**); we therefore set *σ*_*t*_ to be 11.3 ms in the subsequent simulations.

**Fig 6.**
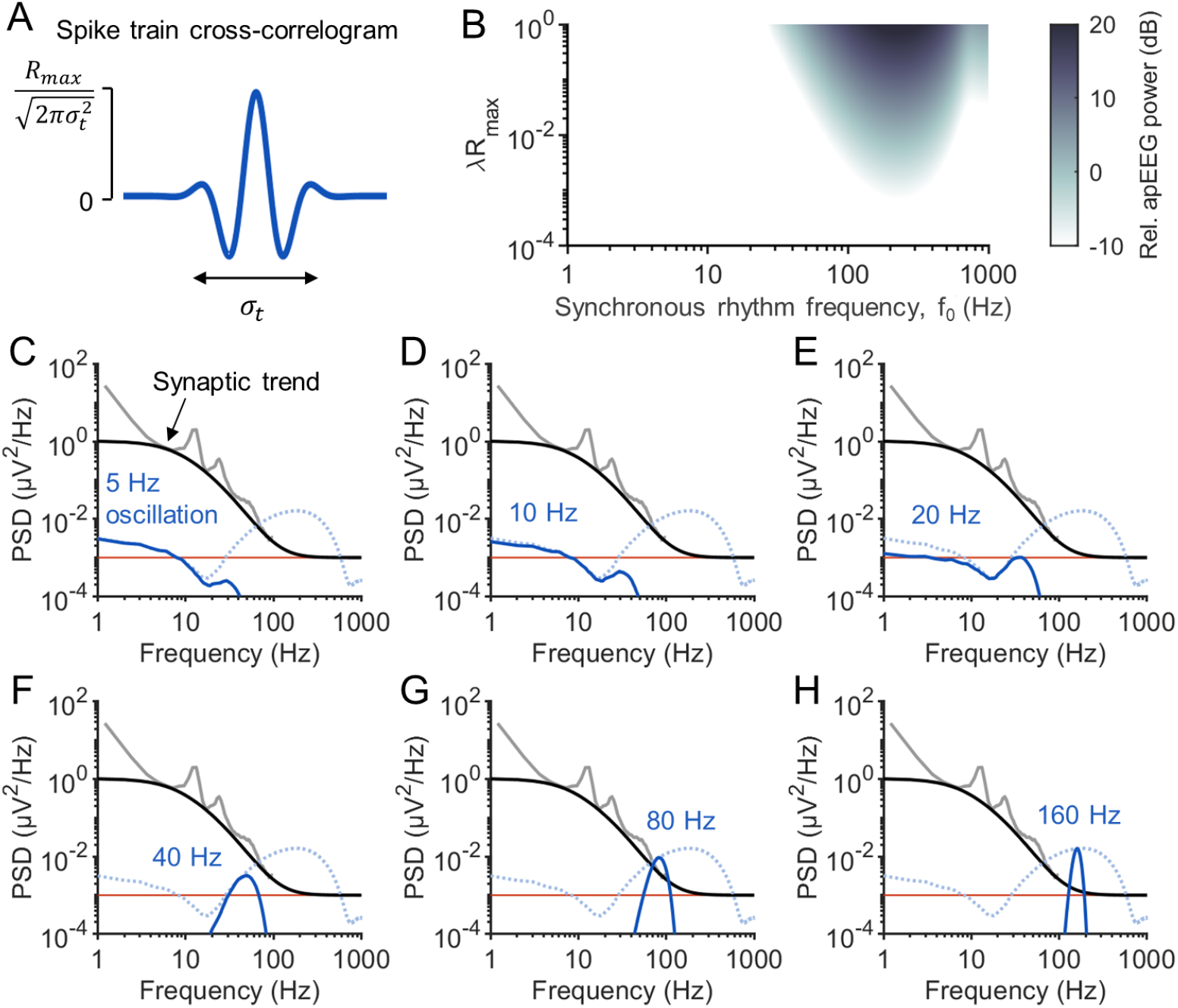
APs can contribute to gamma and higher frequency oscillations. (**A**) The spike train cross-correlation used to model rhythmic spike synchrony, with *σ*_*t*_ = 11.3 ms and *f*_0_ = 40 Hz as an example. (**B**) Simulated amplitudes of spectral peaks generated by APs synchronized at rhythm frequencies between 1 and 1000 Hz. The simulated peak amplitude was defined relative to the fitted spectral trend from **Fig. 5B**. (**C-H**) The average spectra of paralyzed patients (grey) and the fitted spectral trend (black). Notice how at higher frequencies, the spectral trend is constrained by the noise floor (red). The simulated spectrum of the apEEG signal (solid blue) generated by rhythmic spike synchrony at *f*_0_ = 5 Hz (C), 10 Hz (D), 20 Hz (E), 40 Hz (F), 80 Hz (G) and 160 Hz (H). Dotted blue line: the apEEG spectrum generated by the brain with the same average firing rate and maximal correlation as solid blue lines, but with perfectly synchronized spikes, i.e., *σ*_*t*_ = 0 ms.

To evaluate the relevance of simulated apEEG amplitudes, we computed its power relative to the synaptic trend fit to the data of paralyzed subjects (**Fig. 5B**). In this way, we could compare the oscillatory power generated by APs to a “null” EEG spectrum that exhibited no brain rhythms.

When the firing rate or the magnitude of the spiking correlation was too low, APs could not generate any detectable EEG signals (**Fig. 6B**). Interestingly, when the product of the two scaling factors in our model, *λR*_*max*_, was above approximately 10^*−*3^, synchronous rhythmic AP firing could generate pronounced spectral peaks in the EEG spectrum, but only if the oscillation frequency was around 200 Hz. When the value of *λR*_*max*_ was further increased, the regime of oscillation frequencies that produced detectable apEEG signals expanded (**Fig. 6B**).

To better illustrate how these values arose, we plotted one specific example when *λ* = 1 Hz and *R*_*max*_ = 0.2. By superimposing the apEEG spectra on the example EEG data, one can see that APs with slower synchronous rhythm frequencies, *f*_0_, produce apEEG signals well below the amplitude of the EEG spectral trend (**Fig. 6C-E**). However, at higher *f*_0_, the amplitude of the generated spectral peak increased significantly. The amplitudes of these peaks trace out the spectrum generated by the model when synchrony is entirely aperiodic with zero jitter (**Fig. 4F-H**). It follows that the amplitude of apEEG rhythms can be predicted by the simplified synchrony model described in the previous section.

For an 80 Hz rhythm, the parameter combination *λR*_*max*_ needs to be at least 10^*−*2^ for APs to produce a peak with 1% the amplitude of the background spectral trend, and be at least 10^*−*1^ to produce a signal of equal or greater amplitude than the spectral trend (**Fig. 6B**). These values are commensurate with average firing rates between 0.1 and 1 Hz and maximal correlation values around 0.1, which are within the range of values measured experimentally (see Methods, Determining ranges for parameter values). In contrast, no reasonable set of parameters would allow APs to contribute significantly to EEG signals of lower frequency rhythms. We conclude that lower frequency EEG rhythms likely reflect purely synaptic activity, whereas rhythmic EEG signals in the gamma range or higher may be generated by APs and thus directly reflect spiking activity.

## Discussion

We have shown that asynchronous spiking activity cannot cross the noise floor of EEG amplifiers and that, while synchronous aperiodic spiking can generate detectable EEG signals, these signals are dwarfed by the contributions of synaptic currents. On the other hand, we found that rhythmic spiking activity can potentially generate significant EEG oscillations depending on the frequency of such rhythmicity. Together, our results provide quantitative insights into the neural basis of EEG and have direct practical implications for interpreting EEG spectra.

### Interpreting and analyzing EEG spectra

To detrend or not to detrend EEG spectra depends on the mechanisms underlying the changes in the spectral trend^3^. Our results therefore have several important implications for spectral detrending. First, our analysis shows that the high frequency plateau in EEG spectra is entirely determined by excitatory synaptic currents, muscle activity, and amplifier noise. Consequently, any broadband power beyond the frequency range of excitatory synaptic timescales must come from additive noise processes, and therefore should be removed through subtractive detrending^5^. This point highlights that detrending requires different methodologies at high and low frequencies. While the high frequency plateau should be corrected through subtractive detrending, synaptic timescales still need to be corrected divisively^3^. Notably, “whitening” EEG spectra typically involves subtracting the log slope of the spectrum^1, 49^, which is a divisive operation. Our results suggest that this process will overestimate changes in higher frequency EEG oscillations, particularly above 30 Hz. We illustrate this point using a toy model in **Fig. 7**. Suppose two EEG spectra are being compared, one of which reflects a lower excitatory to inhibitory (E:I) ratio and less electromyogram (EMG) (**Fig. 7A1,B1**). Due to the difference in E:I ratio, the spectra need to be detrended of synaptic timescales prior to comparison, as described previously^3, 4^ (**Fig. 7C1**). However, in cases where the high frequency plateau is also changing, for example due to differences in muscle tone, the high frequency plateau needs to be subtracted first. Otherwise, peak estimates will be nonuniformly biased (**Fig. 7D1-E1**). Importantly, correcting for synaptic timescales assumes that spectral peaks are generated by synaptic currents. Our results suggest that this is may not be the case for high frequency oscillations, because these oscillations may be generated principally by APs. In this case, correcting for synaptic timescales when analyzing high frequency oscillations would lead to incorrect conclusions (**Fig. 7A2-E2**). Our results thus strongly indicate that high frequency oscillations should be analyzed separately from lower frequency oscillations. Our simulations provide a reasonable frequency range where oscillations can be generated by AP activity and therefore suggest principled cutoff frequencies for performing different spectral trend analyses. However, this frequency range depended on parameters that we still lack brain-wide estimates for, specifically levels of spiking synchrony and average firing rates. Spectral analysis of EEG would thus benefit from future experimental work investigating these parameters in more detail.

**Fig 7.**
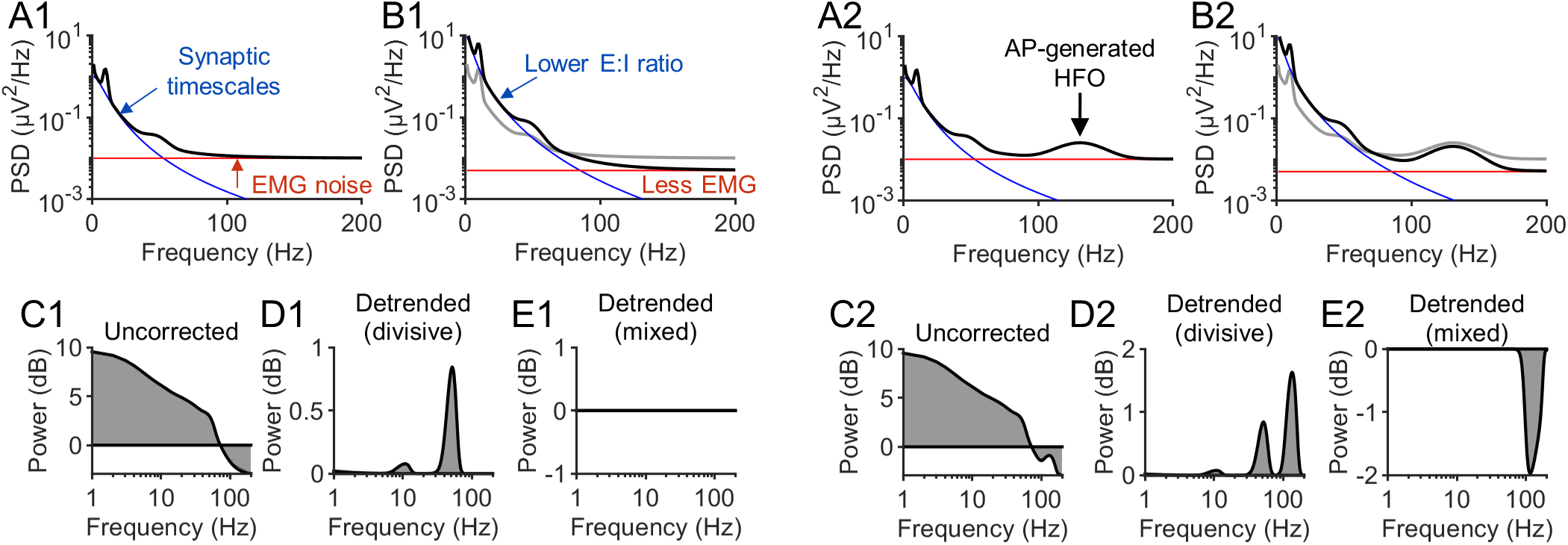
A toy model illustrating the implications of this study’s results on spectral detrending. (**A1**) EEG spectrum (black) modelled with (i) Gaussian functions at 10 and 40 Hz, a constant offset and 1/f noise at low frequencies, all filtered by synaptic timescales (the synaptic timescales fitted in **Fig. 5** were used), plus (ii) additive EMG noise. The equation describing the spectrum was *P* (*f*) = (1 + *α*(*f*) + *γ*(*f*) + 1/*f* ^2^)*P*_*syn*_ + noise, where *α* and *γ* represent the Gaussian peaks. The spectral density from synaptic timescales (blue) and additive noise (red) are overlaid on the simulated spectrum. (**B1**) Same as in A1, but the EMG noise has been decreased by 50% and the E:I ratio was decreased *∼*2.5 fold. The spectrum from panel A1 is shown in grey for comparison. The amplitude of the the Gaussian functions were not changed. (**C1**) Power of spectrum after parameter modifications relative to before (i.e., black vs gray lines in B1). Differences in the spectral trend were not corrected for. (**D1**) The same as in C1, but with the spectral trend, defined as *P*_*syn*_ + noise, removed divisively from the spectra. Notice how this analysis artifactually displays an increase in the alpha and especially the gamma peak, even though these components of the spectrum were not changed. (**E1**) The same as in C1, but with spectra detrended using a mixed approach. Here, the EMG noise was subtracted prior to detrending divisively with the synaptic timescales, producing correctly no changes in rhythmic power. (**A2-E2**) The same as in A1-E2, respectively, but with the inclusion of a high frequency oscillation (HFO) generated by APs using the equation *P* (*f*) = (1 + *α*(*f*) + *γ*(*f*) + 1/*f* ^2^)*P*_*syn*_ + *HFO*(*f*) + noise.

### Neural basis of EEG

Past work has found that apEEG signals can account for up to 20% of EEG rhythms^29^, contrary to our findings. In this previous study, the relative contributions of APs and synaptic currents to the single-neuron dipole were investigated, concluding that APs contribute a large fraction of the single-neuron dipole signal. Our simulations indicated that the average unitary apEEG response is approximately 0.08 pV^2^, whereas the average single-neuron EEG power generated in our passive simulations was 0.09 pV^2^. Thus, at the single-neuron level, our results agree with this previous study, assuming an average firing rate of approximately 0.25 Hz. However, this ratio would only persist in the ensemble EEG if the synaptic and AP components were similarly coherent. Our findings indicate that, even at upper parameter bounds, APs and their after-hyperpolarizations can yield EEG signals with only 1% the spectral density of synaptically-generated EEG signals. We thus conclude that the relative contribution of APs and synaptic currents to single-neuron signals does not translate into ensemble EEG signals.

Our work also has implications for understanding the cellular basis of EEG signals. In particular, EEG signals are typically thought to be generated by pyramidal neurons^24, 50^. However, despite the prototypical morphology of an inhibitory neuron as a stellate cell, with a closed field structure that would prevent them from generating EEG signals^51^, the reconstructed morphologies of the Blue Brain models^34^ displayed significant asymmetries, allowing interneurons to generate apEEG signals only four-fold smaller than excitatory neurons. This result is broadly in line with those of Tenke et al.^52^, who found that small asymmetries in stellate cells allow them to exhibit open field configurations and generate significant current source densities. Moreover, we found that almost the entirety of the apEEG signal amplitude is conferred by spiking synchrony. While cell-type specific synchrony was not included in our modelling, its inclusion would likely boost the relative contributions of inhibitory neurons, as their APs tend to be more highly synchronized than those of excitatory neurons^53^. This observation, combined with the fact that inhibitory neurons tend to fire faster than excitatory neurons and therefore contribute more APs, suggests that inhibitory neurons may contribute significantly to the apEEG signal. In future work, it would be interesting to quantify precisely how much these phenomena compensate for the weaker unitary AP responses and lower abundance of inhibitory neurons. Combined with our other results, these insights may inform what cell types are predominately responsible for high frequency EEG rhythms.

### Modelling assumptions and limitations

In modelling the apEEG signal, we assumed that the AP component of the EEG is independent of the synaptically-generated EEG signal. Although synaptic activity of a population is not statistically independent from its APs, we do not believe that assuming independence distorts our conclusions. Firstly, at the single-cell level, we confirmed that AP power and synaptic power are approximately independent. This can be understood by recognizing that, because of the low firing rate of APs, the majority of power from subthreshold fluctuation will be unrelated to the synaptic events that elicit spikes. Therefore, the more pertinent assumption is that in **Eq. 4** we assume a null cross-spectrum between electric fields generated by APs and postsynaptic potentials (PSPs). We reason that this assumption is plausible because the electric fields generated by APs are determined by the gross morphologies of the presynaptic neurons, whereas the fields generated by PSPs are determined by the precise locations of the synapses on the postsynaptic neurons^3, 54^. Consequently, even if the timing of these events are correlated, the orientation of the resulting current dipoles would only be weakly related on average. The AP-PSP cross-spectrum could be estimated from simulations of neural circuits with morphologically detailed neuron models. However, the results would likely depend strongly on the precise network topology and sub-cellular targeting of synaptic connections which remain only vaguely constrained by experiments. For all the above reasons, we leave these calculations to future studies.

In modelling various neural oscillations, we assumed a constant spatial extent over which spiking activity is coherent. However, lower frequency oscillations are thought to recruit more widely distributed neural ensembles than higher frequency oscillations^55^. Because we did not find reliable parameter values for the spatial extents of each brain rhythm, we assumed that rhythms differed only in their frequency of oscillation. Future work incorporating the spatial topology of different brain rhythms could provide a more complete picture of the role APs play in EEG oscillations.

Finally, recent work has suggested that APs propagating along axons generate dipoles that contribute to LFP recordings^56^. On the other hand, it has been argued that due to the random orientations of axonal termination segments, these signals would not contribute to EEG^29^. In the current study, we only considered back-propagating APs. However, a similar theoretical framework could be used to investigate the electric fields generated by forward propagating APs in more detail. Including these contributions could increase the amplitude of the unitary AP response, potentially expanding the parameter range that permits AP-generated EEG oscillation. On the other hand, including forward propagating APs would not change our finding that jitter in spike timing prohibits high frequency aperiodic apEEG signals, and therefore our conclusions pertaining to the spectral trend would be unaffected.

## Conclusion

Based on our findings, we conclude that APs cannot contribute to the EEG spectral trend, which can be explained entirely by synaptic timescales, electromyogram contamination, and amplifier noise. However, we also conclude that APs can produce narrowband EEG power at high frequencies. These results together suggest that high frequency oscillations and low frequency oscillations interact with the spectral trend differently. While low frequency oscillations require detrending of synaptic timescales, applying this analysis to high frequency oscillations will likely produce incorrect results, and these high frequency oscillations should thus be analyzed separately. Altogether, this work indicates that the interactions between spectral peaks and the spectral trend is frequency dependent, further highlighting that the EEG spectral trend is not a singular phenomenon and should not be removed as a single parameterized function.

## Methods

### Biophysical simulations of unitary AP responses

To calculate the unitary AP responses of various neuron types, we simulated 1034 biophysical neuron models originally developed by the Blue Brain Project^34^. These models have detailed morphological reconstructions of the dendritic arbours and 13 voltage-dependent channels distributed throughout the axonal, somatic, and dendritic segments. For the present work, background synaptic input was added to the model to drive AP firing by distributing 1 excitatory synapse and 0.15 inhibitory synapses per µm of dendrite. The average firing rate of all synapses was set to 1.75 Hz and the ratio of excitation to inhibition, defined as the ratio of mean activation rates between excitatory and inhibitory synapses, was tuned for each neuron model to bring the firing rate above 1 Hz and below 40 Hz. This ensured that APs occurred sparsely enough that the electric fields they generated were independent of one another. To compute each neuron’s unitary AP response, every model was simulated to obtain at least 10 spikes for averaging.

Models were simulated using the python package LFPy^57^, built on top of the NEURON simulation environment^58^, and the single-neuron dipole generated at each time point was calculated using the totality of the current in the dendritic and somatic compartments, as described by Næss et al.^30^. APs were identified in the somatic compartment using MATLAB’s findpeaks algorithm with a minimum peak height set to 0 mV. The spike-triggered average of the single-neuron dipole was then computed for each neuron model.

### Dendrite asymmetry index

Each dendritic arbour was defined by *N* truncated cones with volumes *V*_*i*_ and midpoints ***x***_*i*_, for *i* ∈ {1, 2, …*N*}. The dendrite asymmetry index was then defined as

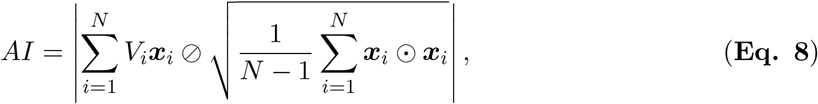

where |·| denotes the Euclidean norm, while ⊘ and ⊙ denote element-wise division and multiplication, respectively. The calculation of this index is illustrated in **Fig. S7**. Conceptually, this equation measures how far the weighted centroid of the dendrites is from zero, normalized in a sense by the span of the dendritic tree. We found that normalization by the actual span of the dendrites over-penalized cells with long apical dendrites. Conversely, without any normalization, cells with large dendritic spans could have relatively symmetric dendrites but large absolute asymmetries, causing their apEEG signals to be overestimated. We found that normalizing by the standard deviation balanced these two extremes well.

### Monte Carlo simulations of ensemble apEEG spectrum

To estimate the ensemble apEEG signal, we evaluated **Eq. 4** numerically using the New York Head model lead field^39^. The New York Head model is based on the ICBM152 v6 brain template^59, 60^, and has the EEG lead field calculated at approximately 75,000 cortical mesh points. All the simulations results here are based on the potential at the Cz electrode site measured against the common average reference.

Evaluating the second term in **Eq. 4** requires computing the cross spectra for all pairwise combinations of cortex coordinates and neuron models (*>* 10^15^ unique pairings), making this step intractable. We therefore used a Monte Carlo sampling approach^61^. Because spike synchrony was assumed to be local in nature with

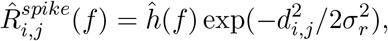

for a spike train cross correlogram of the form *h*(*t*), we could rewrite the second term in **Eq. 4** as follows

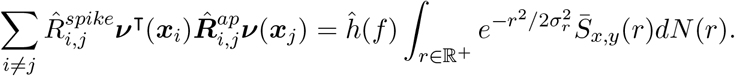

Here, 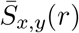 is the expected cross-spectrum between single-neuron EEG signals generated by cells separated by a distance *r*. Note the assumption that *h* is independent of space.

The density *dN* (*r*) reflects the number of neuron pairs in the cortex separated by a distance *r*. We estimated this density as described previously^3^. Briefly, we started by sampling cortex coordinates, ***x***_*i*_, for *i* ∈ {1, 2, …10000}, from the New York head model. Then, for each coordinate, we calculated the total cortical surface area contained in balls of radii *r* ranging from 0 to 200 mm (**Fig. S4A**). Assuming a uniform distribution of 16 billion neurons across the cortical surface area^40^, we determined the empirical density of neuron pairs for each pairwise distance, *r* (**Fig. S4B**). For each pairwise distance, the expected cross-spectrum was computed based on neuron models sampled proportionally to their relative abundance (**Fig. 3B**), and placed at two locations in the cortex; the first location was sampled uniformly and the second location was sampled relative to the first location, according to the density function *dN*.

We terminated our Monte Carlo simulations when there was less than 1% probability that our estimate was off by an absolute error of *δ*_*abs*_ = 4 × 10^*−*25^ µV^2^, which we determined conservatively using Chebyshev’s inequality^61, 62^. This absolute error rate translates into an error in the ensemble EEG spectrum of approximately 10^*−*4^ µV^2^ when all APs are firing in perfect synchrony, an error bound that was chosen because it was equal to the noise floor of high resolution, low noise EEG data^41^. This termination condition was reached after approximately 44 million samples.

### Determining ranges for parameter values

#### Magnitude and timescale of correlation

Co-tuned neurons in area MT exhibit correlations of approximately *R*_*max*_ = 0.2 with a timescale estimated to be around *σ*_*t*_ = 11.3 ms^46^. Unless otherise stated, we used these values in our example simulations. For two reasons, these parameter values likely represent upper bounds for brain-wide spike train correlations. First, neurons that are not co-tuned exhibit less correlated activity^63^, meaning that the average spike synchrony among all neighbouring neurons is likely lower than that between co-tuned neurons. Consistent with this, a survey of studies across different experimental paradigms and cortical areas found correlation values ranging between 0.05 and 0.25^44^. In area MT, the timescale of correlation was found to be around 10 ms^46^, whereas studies in V4 indicate timescales of tens or hundreds of milliseconds^43, 64^. We therefore took a liberal range of 10 to 100 ms as an acceptable range for *σ*_*t*_.

#### Mean firing rate

Because activity is sparse in the cortex, most neurons are silent at any particular moment; this silent fraction has been estimated to be up to 90% of all cells^65^. As a consequence, the average firing rate across the entire brain is significantly lower than would be expected from measurements of only responsive neurons, with estimates averaging around 0.1 to 2 Hz^65–67^.

### EEG data and spectral trend fitting

The experimental data shown in **Figs. 3E, 4C** and **5** were extracted directly from the figures in Scheer et al.^41^ and Whitham et al.^20^. The spectra from Whitham et al.^20^ were fit with the equation

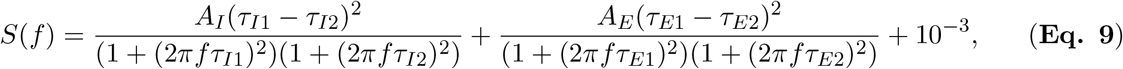

where *τ*_*I*1_ and *τ*_*I*2_ are the rise and decay time constants associated with inhibitory synaptic responses and *τ*_*E*1_ and *τ*_*E*2_ are the rise and decay time constants associated with excitatory synaptic responses, while *A*_*I*_ and *A*_*E*_ govern the relative contribution of inhibitory and excitatory currents to the EEG spectrum. This equation was adapted from Eq. 6 in Brake et al.^3^, except that here the high frequency plateau was explicitly decomposed into excitatory synaptic contributions and the noise floor, which was empirically determined to be *∼*10^*−*3^ µV^2^Hz^*−*1^. The same *τ* values were used to fit both the unparalyzed and paralyzed data; only the scaling factors *A*_*I*_ and *A*_*E*_ were adjusted.

## Code availability

Code used to run simulations, analyze data, and generate manuscript figures is available on GitHub (github.com/niklasbrake/apEEG modelling).

## Acknowledgements

This work was supported by the Natural Sciences and Engineering Research Council of Canada (NSERC) discovery grant (RGPIN-2019-04520) to AK. NB was supported by the NSERC-CREATE in Complex Dynamics Graduate Scholarship and the Fonds de recherche du Québec – Nature et technologies (FRQNT) doctoral training scholarship. The funders had no role in study design, data collection and analysis, decision to publish, or preparation of the manuscript. This research was also enabled in part by support provided by Calcul Québec (calculquebec.ca) and the Digital Research Alliance of Canada (alliancecan.ca).

## Author contributions

**Niklas Brake**: Conceptualization, Methodology, Investigation, Writing-Original draft preparation, Visualization. **Anmar Khadra**: Supervision, Funding Acquisition, Writing-Reviewing and Editing.

## Supporting Information

**Supplementary Fig. S1.**
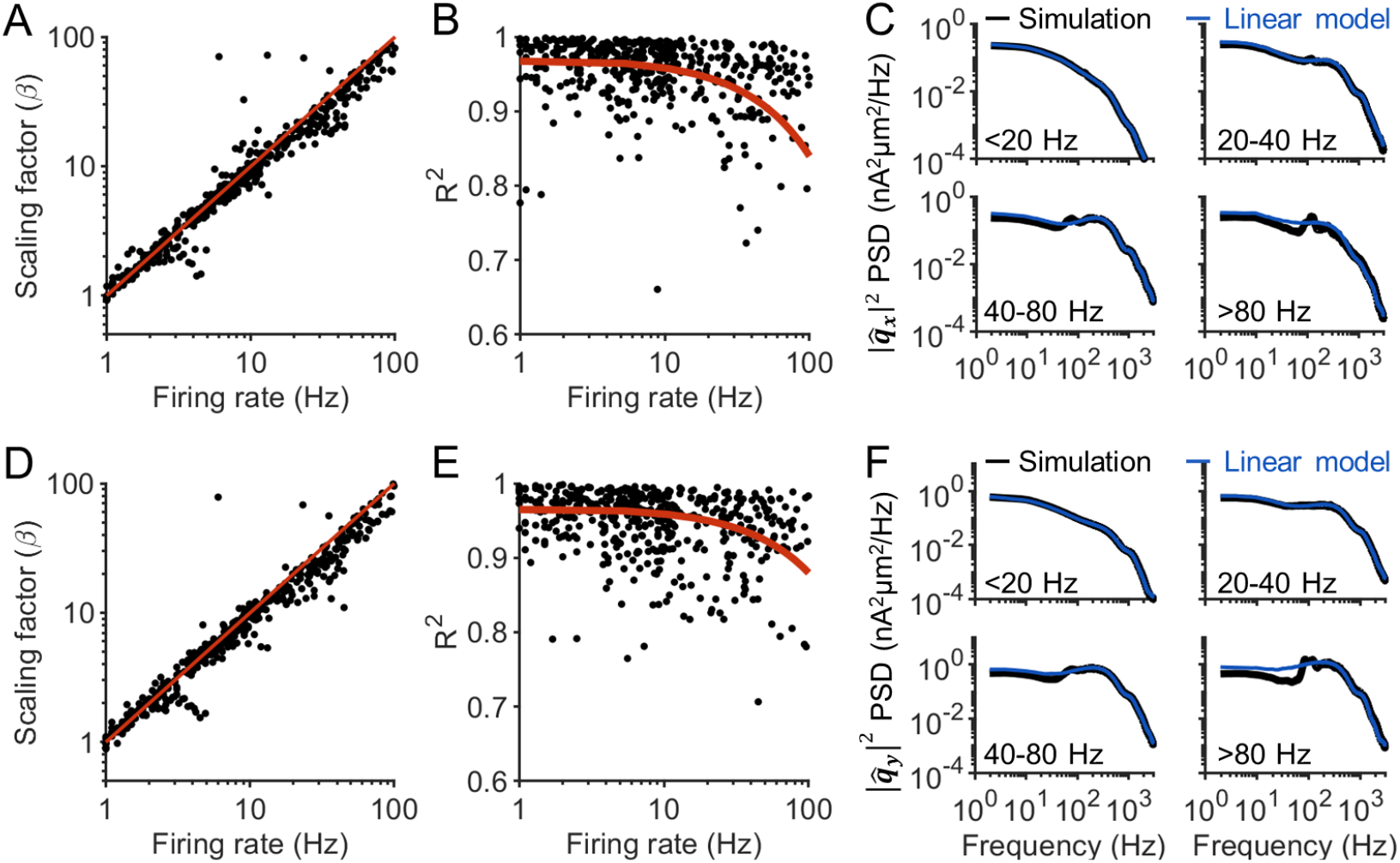
Scaling of the x and y components of the unitary AP spectrum. (**A-C**) Same as **Fig. 2**, but for the *x* component of the single-neuron dipole. (**D-F**) Same as **Fig. 2**, but for the *y* component of the single-neuron dipole.

**Supplementary Fig. S2.**
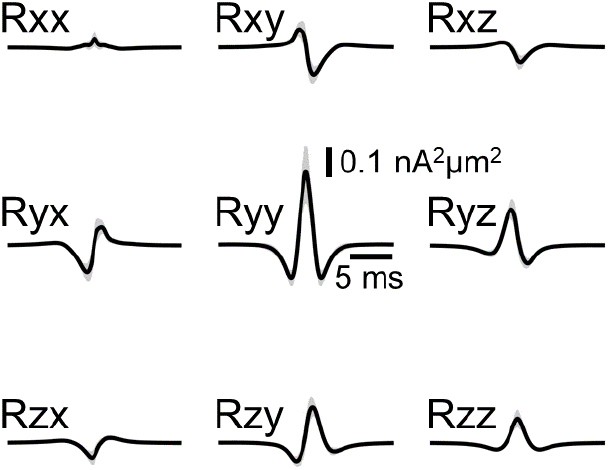
Cross-correlation among unitary AP responses. Solid black line indicates average across all pairs of 1035 neuron models, weighted by the relative abundance of the pairing (**Fig. 3B**). Shading reflects 95% confidence interval of the mean.

**Supplementary Fig. S3.**
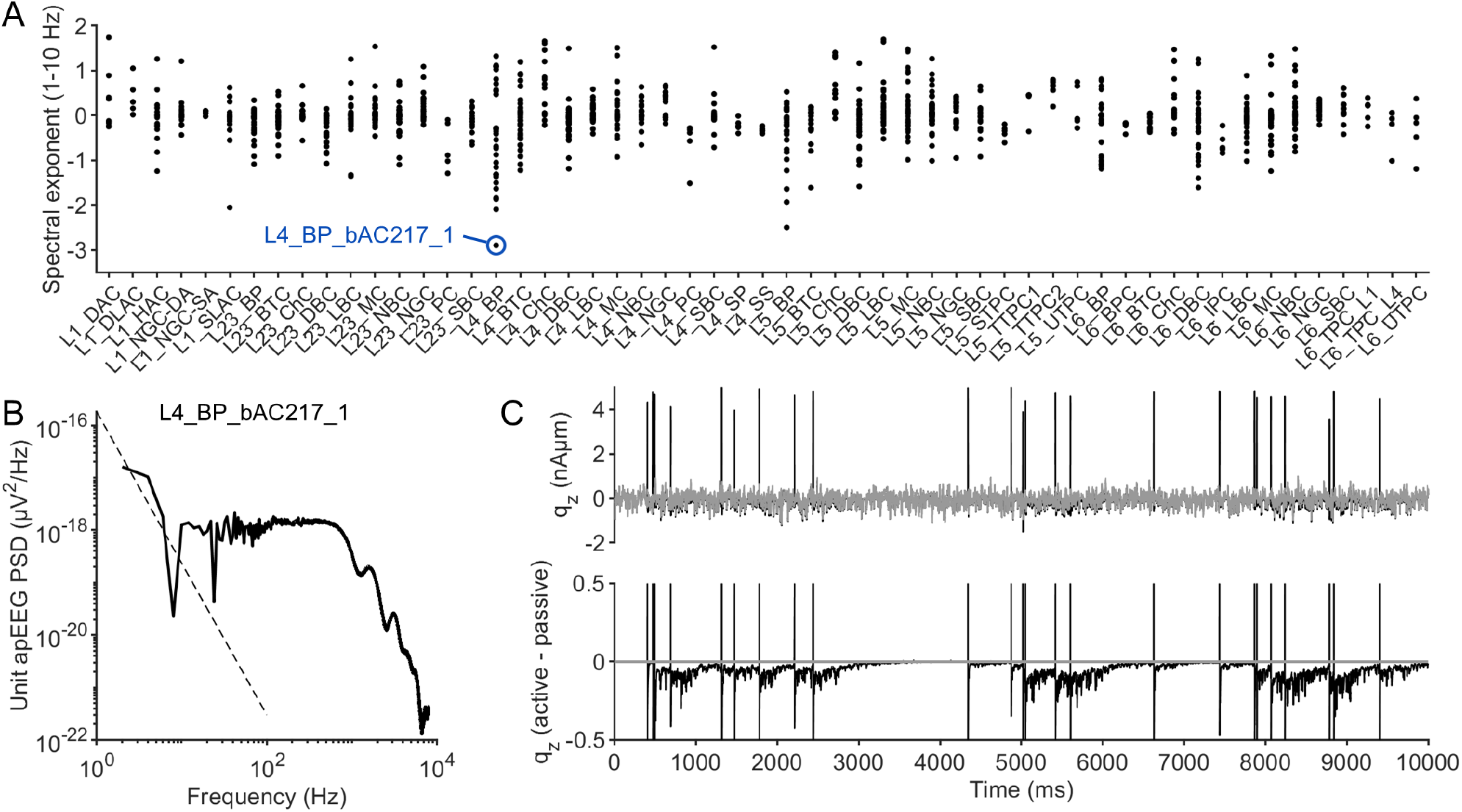
Low frequency apEEG power generated by afterpotentials. (**A**) The slope of the unitary AP spectrum for every neuron model, calculated between 1-10 Hz. A neuron (ID: L4 BP bAC217 1) with a particular negative slope is indicated in blue. (**B**) Power spectrum of the unitary AP spectrum for the neuron indicated in panel A, with 1/f trend fitted at low frequencies (dashed black line). (**C**) Top: z component of the single-neuron dipole of the neuron indicated in panel A (black) and with somatic and axonal sodium channels removed to generate a passive model (grey). Bottom: Difference between the active and passive neuron models. Note that the spikes have been truncated at *±*0.5 nA µm. After each spike, the active model’s dipole takes hundreds of milliseconds to reconverge to the passive model. The same phenomena were observed in the dipoles x and y components.

**Supplementary Fig. S4.**
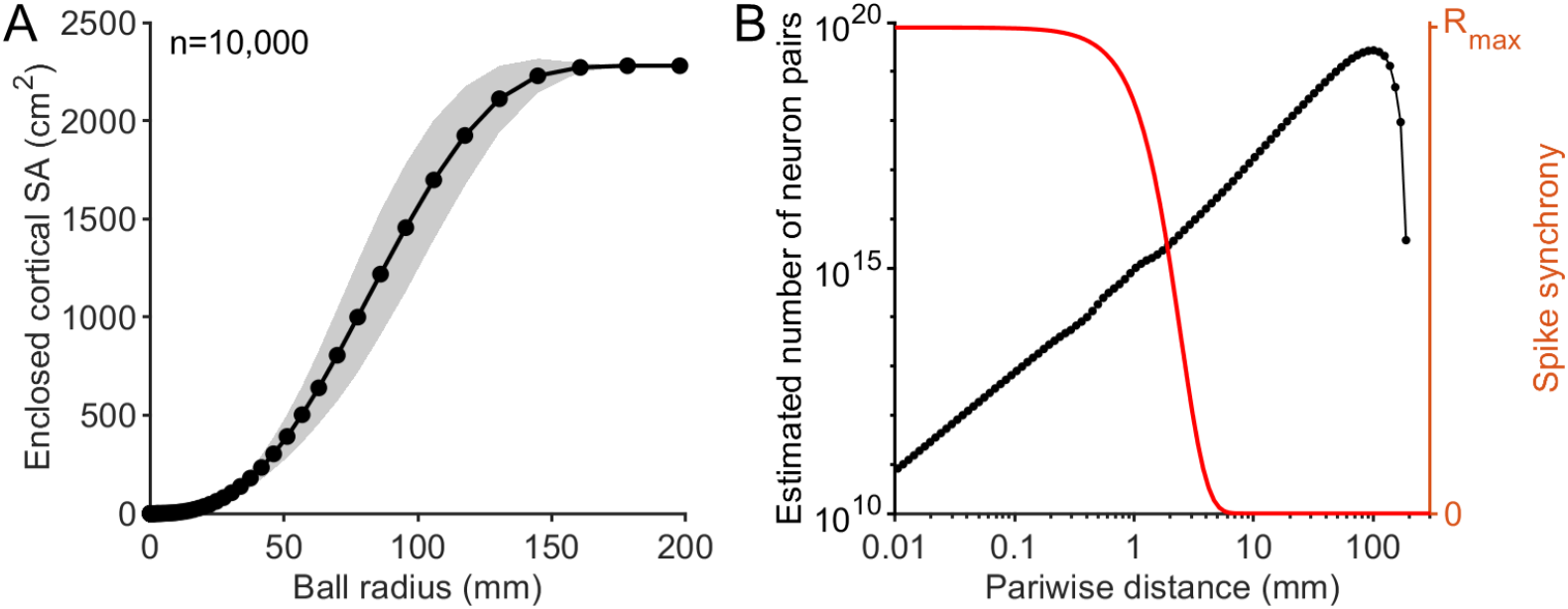
Distribution of neuron pairs with respect to pairwise distance. (**A**) Surface area (SA) of the cortex from New York head model enclosed within balls of increasing radii, with the origin of the ball placed at 10,000 cortical locations. Black dots indicate the discrete ball radii for which the surface area was calculated. Shading reflects standard deviation across the 10,000 starting points. (**B**) The derivative of the surface area with respect to radius (black), scaled to obtain the density of neuron pairs for each pairwise distance (see Methods). The red curve illustrates the coupling kernel, as in **Fig. 4A**. The vast majority of neuron pairs are separated by more than 10 mm and are therefore not correlated in the model.

**Supplementary Fig. S5.**
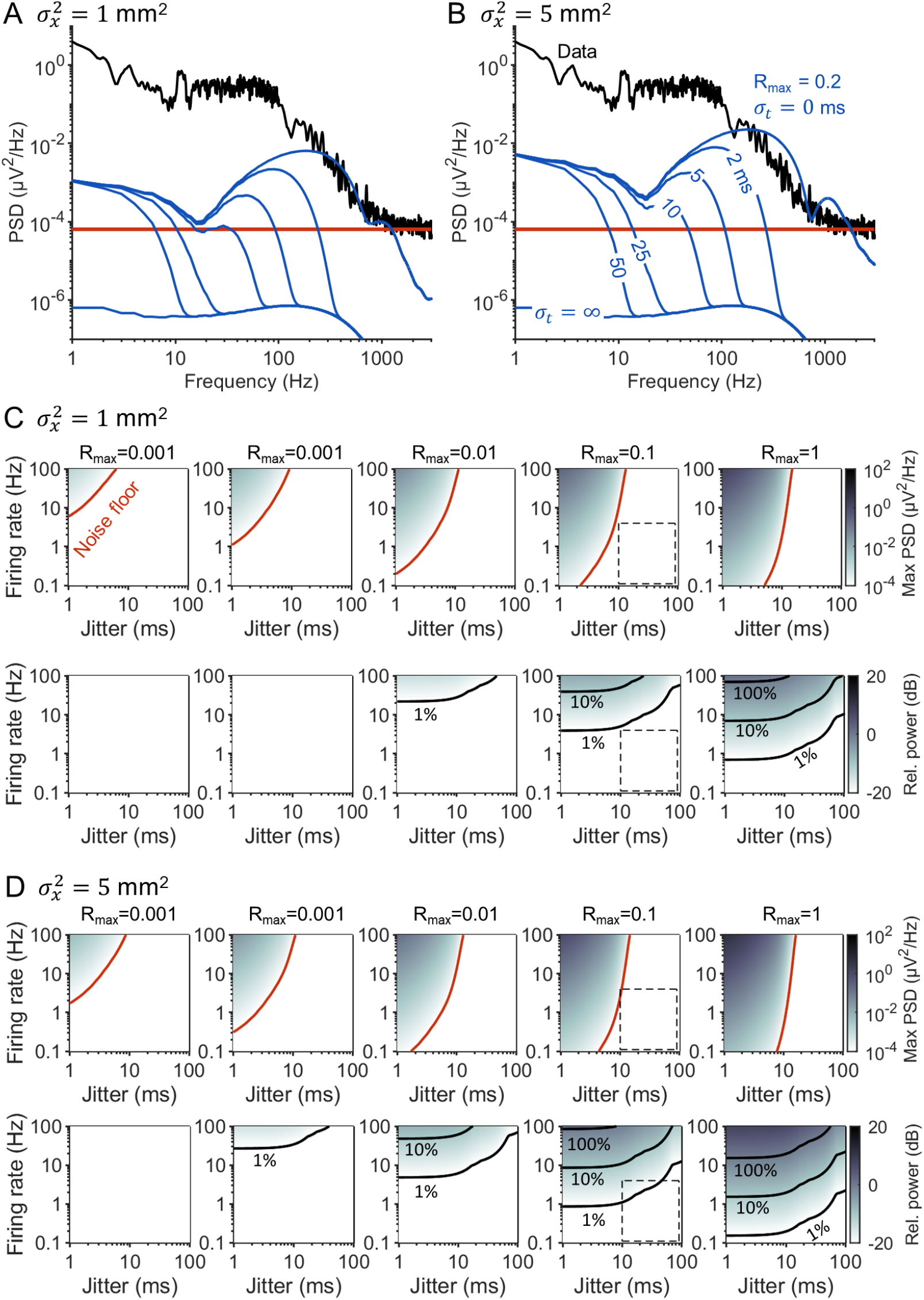
Sensitivity of aperiodic apEEG to *σ*_*x*_. (**A**) Same as in **Fig. 4B**, but for 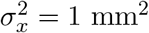. (**B**) Same as in **Fig. 4B**, but for 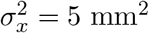. (**C**) Same as in **Fig. 4C** (top) and **Fig. 4D** (bottom), but for 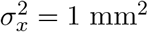. (**D**) Same as in **Fig. 4C** (top) and **Fig. 4D** (bottom), but for 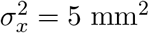.

**Supplementary Fig. S6.**
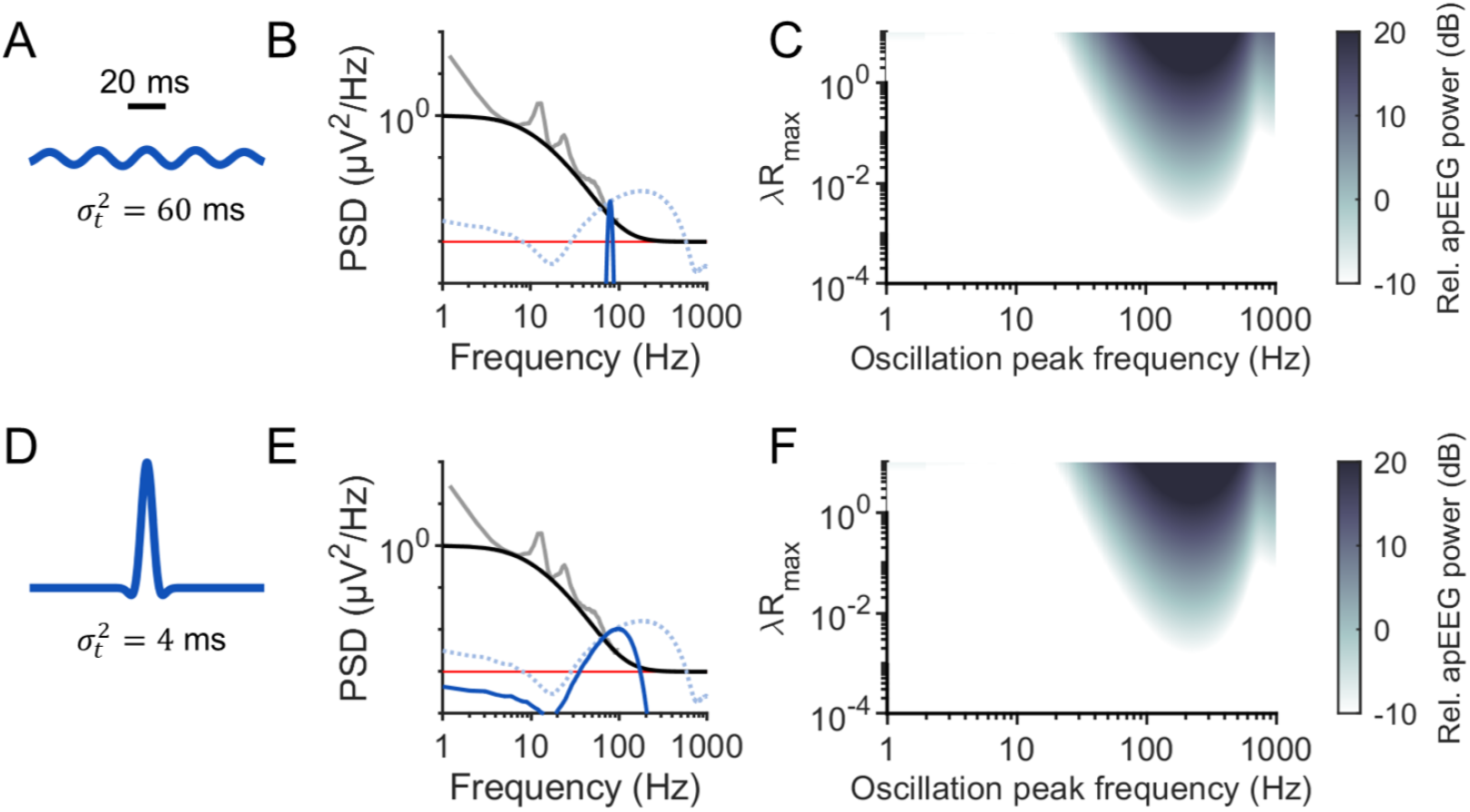
Timescale of correlation affects spectral peak width of apEEG rhythm. (**A**) Plot of **Eq. 7** for 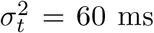 (**B**) Same as in **Fig. 6G**, but for 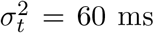. (**C**) Same as in **Fig. 6B**, but for 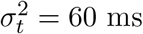. (**D-F**) Same as in A-B, but for 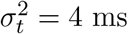 ms.

**Supplementary Fig. S7.**
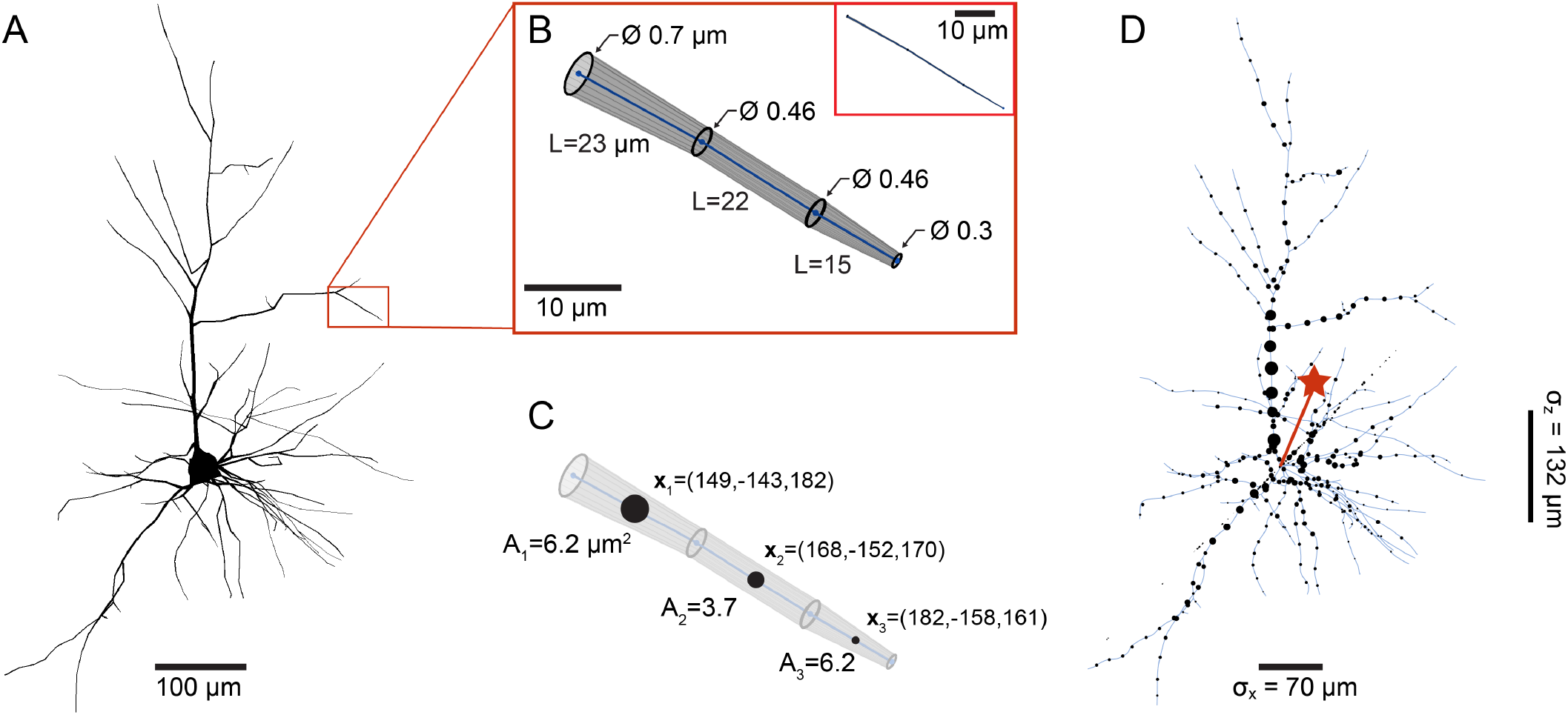
Schematic of dendrite asymmetry index calculation. (**A**) Example morphology of a layer 6 pyramidal cell. Note that the diameter of the dendrites have been increased by a factor of two in the figure to better illustrate the variation in dendrite diameter throughout the arbour. (**B**) Zoomed in view of the indicated dendritic branch, showing that the dendrite morphology is represented by truncated cone segments.The diameter and length of each truncated cone is printed. For illustrative purposes, the dendrite diameter is drawn with a scaling factor of 10. The same dendrite segment with correct proportions is shown in the insert for comparison. (**C**) To calculate the dendrite asymmetry index (**Eq. 8**), each segment is represented by its midpoint in space (***x***_*i*_) and its total volume (*V*_*i*_), calculated as 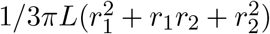. The black dots plotted at the midpoint of each dendrite segment are scaled proportionally to the segment’s volume. (**D**) The black dots represent the midpoints and their sizes represent the volume of all dendrite segments. The red star indicates the result of the asymmetry index calculation (**Eq. 8**), prior to taking the Euclidean norm. The length of the red line is thus the asymmetry index of this neuron. For illustrative purposes, the equation result has been scaled here by 0.01 as otherwise the vector would be too long to depict. Note, however, that the regression in **Fig. 3** holds for any arbitrary scaling of **Eq. 8**.

